# Population Models with Phenotypic Diversity and Fluctuating Environment Illustrate Parameter Regimes of Adaptation vs. Extinction

**DOI:** 10.1101/2025.01.01.631012

**Authors:** William H. Press

## Abstract

A species may survive long-term environmental fluctuations by maintaining a sufficient phenotypic variability to span the changes. Bet-hedging and adaptive mutatability (or mutator strains) are among possible mechanisms. We construct simple population dynamics models for these strategies in a unified framework, with a stochastic environment model and a specifiable environmental carrying capacity, identifying different parameter regimes: Quantitatively, how much hedging is needed or what mutation rate is required? How much selection burden re wild-type is tolerable for the strategy to work? Is the diverse population susceptible to invasion by a reverting wild type better adapted in the short run, but producing an extinction outcome in the long run? We give answers that are quantitative in the context of the specific idealized models, but that can also be qualitatively illuminating to more realistic cases.

## 1 Introduction

We consider how a species may survive, or become extinct, in a fluctuating long-term environment, for example, pre-anthropogenic climate variation. As the external environment changes, a previously well-adapted phenotype can find itself maladapted, leading to extinction of the species. Long understood by ecologists and evolution theorists [1, 2, 3] is that phenotypic diversity can rescue a species from such an outcome if variants adapted to a range of environments are maintained in the gene pool. The problem is how to maintain such diversity in the presence of selection against a maladapted variant—even when that variant could be essential in the future. Evolution does not look or plan ahead.

Among possible mechanisms for maintaining diversity, two are bet hedging [2, 4], the strategy of producing in each generation a mixture of phenotypes, regardless of their relative present fitness; and adaptive mutation or mutability (and the existence of so-called mutator strains) [5, 6, 7], the idea that mutability is a trait itself subject to selection [8, 9], thus able to evolve to greater mutation rates, perhaps conditionally on only certain genes [10, 11, 12], thereby maintaining the required level of gene pool diversity.

Both of these concepts have been at times controversial [13, 14], because both seem to violate the no-look-ahead principle. Bet hedging and adaptive mutation each impose current selection penalties for future selection benefits. In that case, the argument goes, how can either mechanism be maintained in the long term in the face of positive selection for a mutation that disables that mechanism and its cost? Evidently the answer must be a matter of quantitative magnitudes (how small is the cost?) and timescales (how soon does the genetic diversity pay off?) [15]. Today, little remains controversial [16, 17, 18].

This paper constructs population models, small sets of ordinary differential equations (ODEs), that can exemplify the quantitative trade-offs involved in either bet hedging and adaptive mutation. While the models are highly simplified and ignore many real complexities, they may shed light on the quantitative relationships among various parameters that are necessary for successful continuing adaptation.

This is hardly unexplored territory. These issues broadly are the subject of hundreds of papers over the last fifty years. Ref. [17] summarizes the literature pre-2002. Among many more recent examples of work exploring these issues via population models are [19, 20, 21]. What this paper attempts to add to this large literature is a well-defined model architecture that facilitates the quantitative comparison of different mechanisms. The architecture incorporates the constraint of environmental carrying capacity, and it tests the population dynamics against a representation of environmental change that varies stochastically in a well-defined way, as next described.

### 1.1 Environment Interaction Model

The effective population size of a particular phenotype *x*, denoted *n*_*x*_, will be modeled by an ODE of the general conceptual form

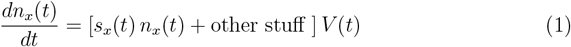

Here *s*_*x*_(*t*) is the intrinsic growth rate, the theoretical rate of population growth absent other limiting factors. A necessary condition for a species to survive is that it have, at least sometimes, a phenotype with a positive value *s*_*x*_, since, otherwise, its population always decays. In equation (1), the long-term environment enters through the time dependence of *s*_*x*_(*t*), which will fluctuate between positive for a well-adapted phenotype to negative for a maladapted one.

We take the time dependence of *s*_*x*_(*t*) to be a bandwidth-limited Gaussian process *G*(*t*) parameterized by a timescale *T*_env_, the “environment timescale.” Specifically,

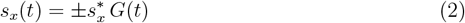

where the constant 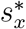 is the coefficient of phenotype *x*’s response to the environment, and the sign ± distinguishes the two genotypes: positive for the well-adapted, negative for the maladapted. *G*(*t*) is specified by its Fourier spectrum,

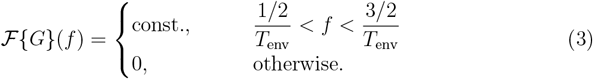

where *ℱ* denotes Fourier transform. Equation (3) says that *G*(*t*) varies with an oscillation period of about *T*_env_, but not exactly periodically. The constant is chosen to give *G*(*t*) unit r.m.s. amplitude. Figure 1 shows realizations of *G*(*t*) for different values of *T*_env_.

**Figure 1:**
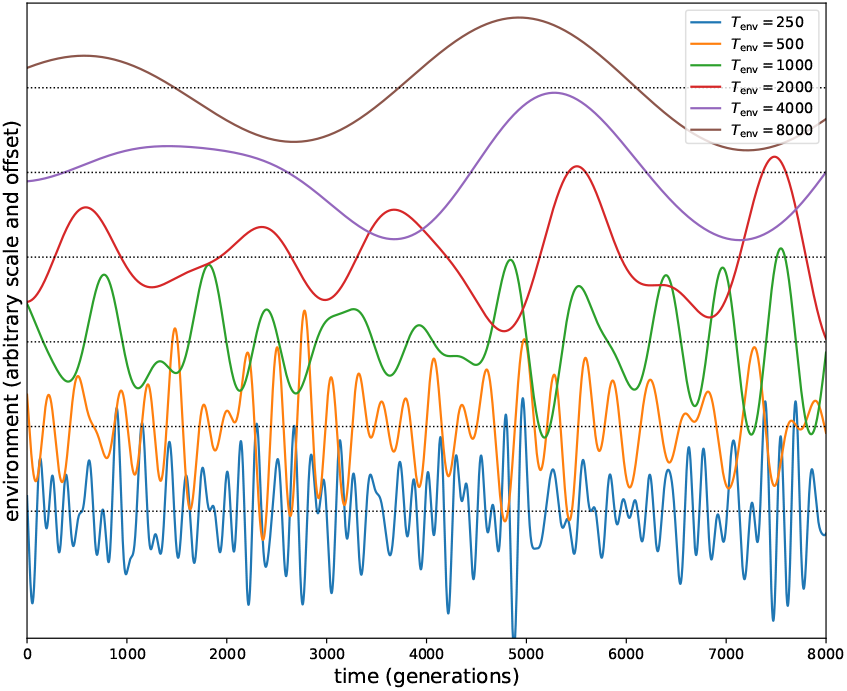
Environment (e.g., climate) is modeled as an aperiodic Gaussian process with a specified characteristic timescale *T*_env_. Shown here are typical realizations with different timescales (arbitrary vertical scale).

For definiteness, we set 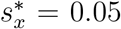. Taken literally, this implies an intrinsic growth rate that yields a net reproductive rate *R*_0_ of *∼* 1.05 for a well-adapted phenotype, *∼* 0.95 for a maladapted one. However, the value of *s*^*∗*^ simply scales the units of time *t*. The adopted value 0.05 makes the figures below more interpretable, but (via scaling) does not limit the models’ generality.

The factor *V* (*t*) in equation (1) is used to enforce the species carrying capacity, a maximum population size that the environment can sustain. If the carrying capacity is *N*_*∗*_, a conceptual specification of *V* is

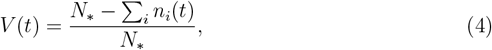

making the growth rate of of all phenotypes *n*_*i*_ of a species taper linearly to zero as the sum of their numbers approaches the species carrying capacity. Some modifications of this will be discussed. (In the textbook literature, *N*_*∗*_ is often denoted *K*.)

### 1.2 Fixation and Loss Timescales

In simple Wright-Fisher theory, a novel allele with positive selection coefficient *s* will fix in a population of effective size *N* with probability

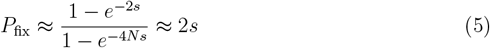

where the second approximate equality holds for *s* ≲ 1, *Ns*≫ 1, which we will generally assume. The characteristic time to fixation is then [22],

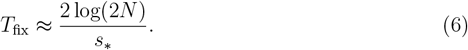

This can be understood as the time for exponential growth through log(2*N*) e-folds at a rate of *s*_*∗*_ e-folds per time. Heuristically, the factor 2 inside the log reflects diploidy, while the factor 2 outside reflects saturation of the exponential increase (S-shaped curve). By similar reasoning, with a minor additive adjustment of the logarithm factor, *T*_fix_ is also the time for an allele with negative *s* to become lost. For simplicity, we take equation (6) as approximating both cases. Likewise, we will not distinguish among related values of *N* inside the logarithm that produce only small fractional changes in *T*_fix_, e.g., distinguish among *N*_*∗*_ (the carrying capacity), *N* (the actual population, when within an order of magnitude of the carrying capacity), or *n*_*x*_ (the population of one allele *x* if it is significant in the population).

## 2 The Scenarios

The several models in §3, below, are defined by sets of ODEs. However, these—and the overall scope of this paper—will be clearer if we first describe the several scenarios that the models represent. The details of each scenario are fanciful, but illustrative of the models’ variables and parameters.

[Scenario 0 (common to all models).] Puffs are small, furry, fictitious animals that live in burrows under the tree roots in mid-latitude forests. A single gene with two alleles regulates the length of their fur. Short-haired puffs thrive in warm climatic periods; long-haired puffs in cold; each does poorly in the opposite circumstance. Burrow space is limited, so, inclusive of both phenotypes, there is a finite ecological carrying capacity, implying a maximum population size.

A minimal set of parameters for Scenario 0 consists of the environmental timescale *T*_env_ (equation 3), the intrinsic growth rate *s*_*∗*_ (equation 2), and the carrying capacity *N* ^*∗*^ (cf. equation 4), the latter two implying a characteristic fixation time *T*_fix_ (equation 6).

### 2.1 Bet Hedging Scenarios

Bet hedging is introduced in Scenario 1:

[Scenario 1 (bet-hedging with no further penalty).] In each generation of puffs of each phenotype, a fraction of the next generation is born with the other phenotype, but this feature otherwise creates no new selection penalty.

Two new parameters, *η*_1_ and *η*_2_ are hereby introduced, the fractions of the opposite (i.e., non-parental) phenotype born in each new generation of short-or long-haired puffs. One may have *η*_1_ = *η*_2_, or else not.

Scenario 1 is not a complete evolutionary picture. Since the ability to produce population diversity is itself a trait that can be lost by mutation, a population with that trait competes against one in which the trait has been lost, reverting it to the (unhedged) wild type. This defines:

[Scenario 2 (evolutionary stability of hedging trait).] The puffs all still have the gene whose alleles are long- and short-hair. Some individuals have the gene (or more complicated pathway) that creates bet-hedging population diversity, but other individuals have reverted to wild type (without hedging). There are now four phenotypes: long/short hair (to be denoted 1 and 2) and with/without bet hedging (to be denoted *a* and *b*).

Scenario 2 adds no new parameters to Scenario 1, other than choosing the starting population sizes of each of the four phenotypes, which (below) will prove irrelevant to the long-term behavior of the model.

It remains a possibility that the genetic apparatus necessary to produce bet hedging imposes, via some unrelated side effect, its own selectivity cost. This might increase the likelihood of reversion to the unhedged (and therefore less survivable) wild type.

[Scenario 3 (added penalty for hedging trait).] Same as Scenario 2, but there is an additional fitness penalty imposed on both the short- and long-haired puffs that carry the bet hedging trait.

The penalty is a new parameter *b*, taken to be the decrement in selection coefficient.

### 2.2 Adaptive Mutation Scenarios

We turn next to scenarios with adaptive mutation.

In Wright-Fisher or similar models, most mutations, even those with positive selection coefficients, do not survive in the population, as already indicated in equation (5), above. Heuristically, this is because growth of numbers in the population is a diffusion process with an absorbing boundary condition at zero individuals, combined with the fact that any one-dimensional random walk is very likely to have zero-crossings.

[Scenario 4 (fixed mutation rate with no penalty).] Baseline Scenario 0, except that a mutation from short- to long-haired puff, or vice versa, occurs randomly at a mean rate *µ* per individual per generation.

For simplicity we will take the rates in each direction as the same, though this assumption would be easy to relax. We refer to a mutation as “effectual” if it has *s >* 0 and survives a draw from the probability in equation (5). Effectual mutations are, in other words, those that reach the continuum regime and can then be modeled by deterministic ODEs rather than stochastic diffusion equations [22]. Because *s* = *s*(*t*) is time varying and can change sign, not all effectual mutations will actually fix; but this is captured by the evolution of the ODEs. Scenario 4 adds a new parameter, which we can take to be the average time in generations between effectual mutations in a population of size *N* ^*∗*^ with the intrinsic growth rate *s* = *s*_*∗*_,

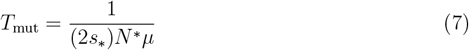

where the factor 2*s*_*∗*_ is from equation (5). For any particular phenotype with population *n < N* ^*∗*^, the average time between effectual mutations is larger, because the population is smaller, by a factor *N* ^*∗*^*/n*; we will model this by generating Poisson random mutations at the rate of equation (7) but accepting them for a phenotype only with probability *n/N* ^*∗*^.

## 3 Model Equations and Behaviors

### 3.1 Baseline Model

Translating Scenario 0 to equations yields,

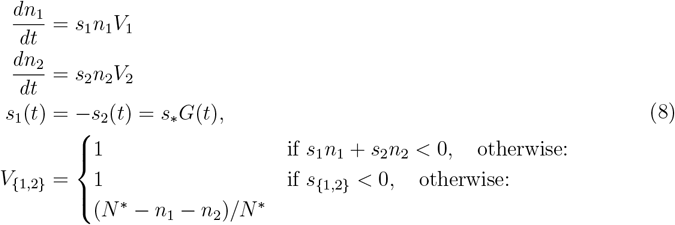

The effective population sizes of short- and long-haired puffs are here *n*_1_ and *n*_2_. The intrinsic growth rates *s*_1_(*t*) and *s*_2_(*t*) are equal in magnitude, opposite in sign, and fluctuate according to the Gaussian process *G*(*t*) of equation (3).

The several cases for carrying capacity factors *V*_1_ and *V*_2_ require some explanation. The basic idea is that a shrinking total species or individual phenotype population should have *V* = 1 (no inhibition of the rate of shrinking), because there is an implied excess of supportive capacity (i.e., burrows); but a phenotype whose increasing number requires new burrows should be limited in the manner of equation (4), above, so that the total population of the species does not exceed the environment carrying capacity *N* ^*∗*^. The intent is to penalize a species’ expanding into an ecological niche with finite carrying capacity, but not to mitigate a contraction due to negative selection.

Although population sizes *N* ^*∗*^, *n*_1_, and *n*_2_ are intuitively defined, the equations simplify with scaling to *N* ^*∗*^ (assumed constant) and defining fractional niche occupation numbers *f*_*i*_ *≡ n*_*i*_*/N* ^*∗*^. Equation (8) then becomes,

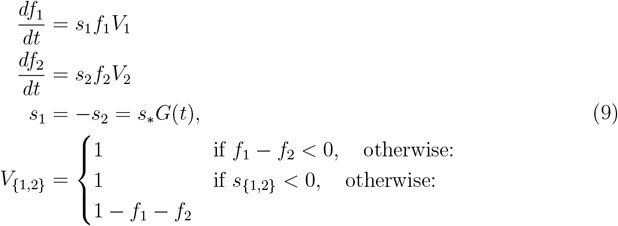

For generic initial conditions *f*_1_, *f*_2_ ≲ 1, *f*_1_ +*f*_2_ ≤ 1, the generic behavior of equation (9) is determined by a single dimensionless parameter, the ratio of the two timescales *T*_env_ (cf. equation 3) and *T*_fix_ (equation 6), expressed in terms of model quantities as

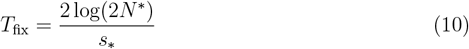

Figure 2 shows a typical case for *T*_env_*/T*_fix_ ≈ 0.5 *<* 1. One sees that the rapidly changing climate allows both long- and short-haired puffs to survive indefinitely in the population, or at least until a statistically unlikely prolongation of warm or cold climate (not shown in the figure). Notably, neither phenotype has time, on average to grow to the environmental carrying capacity, or even nearly so.

**Figure 2:**
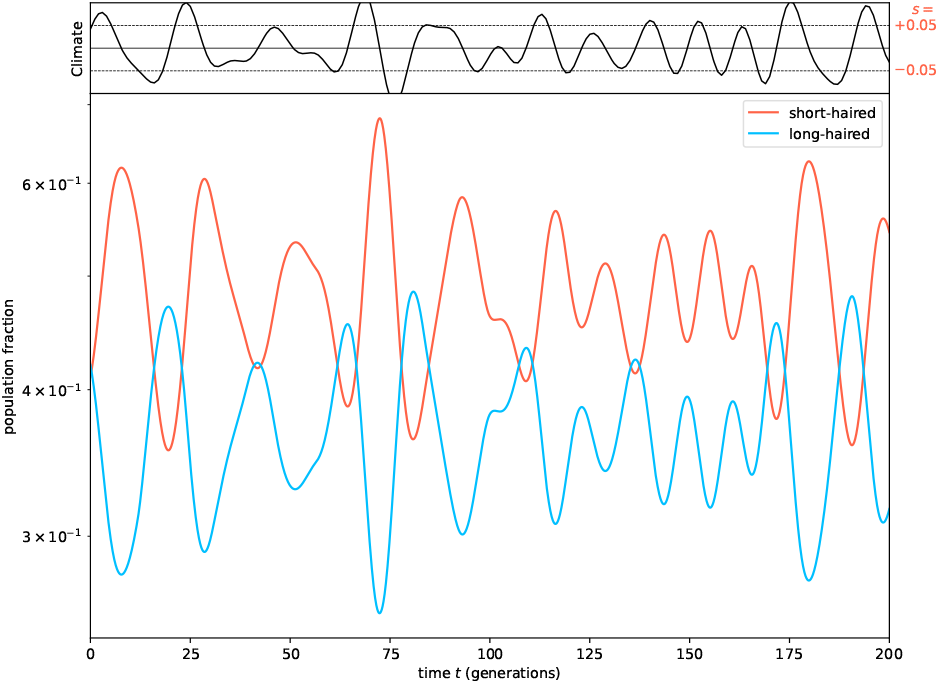
(Scenario 0.) Upper plot: A stochastic realization of environmental fluctuations. Positive (negative) values favor short-(long-) haired puffs. Lower plot: Population dynamics of the two phenotypes, one adapted to hot, the other to cold environments. Here, with *T*_env_*T*_fix_ ≲ 1, neither goes extinct before the changing climate causes it to be favored. But, correspondingly, neither has time to expand to the environmental carrying capacity (here scaled to 1).

Contrast the above to Figure 3 where *T*_env_*/T*_fix_ ≈ 2.0 *>* 1. In this particular realization, both phenotypes survive for a few climate cycles. However, in the lengthy warm period that starts at about generation 550, the long-haired phenotype goes extinct (at around generation 720). The short-haired phenotype increases to the carrying capacity, only to then go extinct when the climate changes to cold.

**Figure 3:**
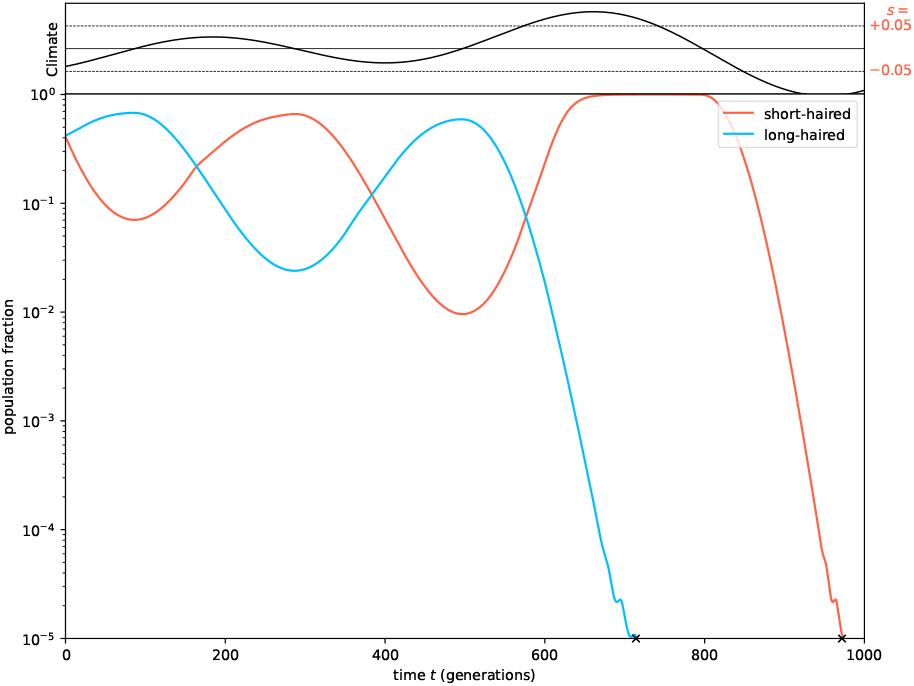
(Scenario 0.) Same as Figure 2, except that now *T*_env_*T*_fix_ *>* 1. Both phenotypes survive a few climate cycles, but eventually each becomes extinct (denoted by X on the time axis) in a prolonged unfavorable climate.

Although the details may differ stochastically (number of cycles before extinction, equilibrium population relative to carrying capacity, etc.) the two behaviors seen in Figures 2 and 3 are the unique generic behaviors of this model, depending on the single parameter *T*_env_*/T*_fix_.

### 3.2 Bet Hedging Models

#### 3.2.1 Bet Hedging with No Further Penalty

The equations for Scenario 1 are

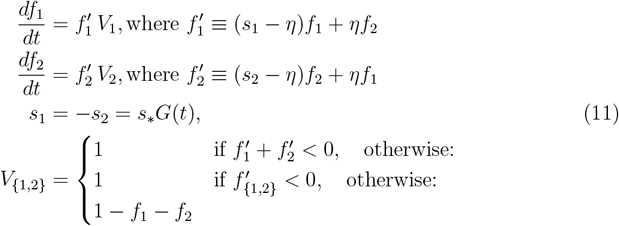

In a rapidly changing environment *T*_env_*/T*_fix_ ≪ 1, Scenario 1 is hardly different from Scenario 0 (Figure 2): Neither phenotype becomes endangered before the climate fluctuates back to its favorability. The species survives. In the opposite case, *T*_env_*/T*_fix_ ≫ 1, the model has only a single generic behavior, that shown in Figure 4. In its favored periods, each phenotype grows to nearly the species carrying capacity, while the other phenotype falls, but only as far as a stable equilibrium value. The equilibrium population of the unfavored phenotype, *f*_*u*_, can be approximately calculated by solving simultaneously *f*_1_ + *f*_2_ *≈* 1 and 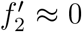, taking *s*_1_ and *s*_2_ as their r.m.s. values *s*_1_ = *−s*_2_ = *±s*_*∗*_. The result is

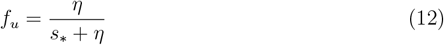

**Figure 4:**
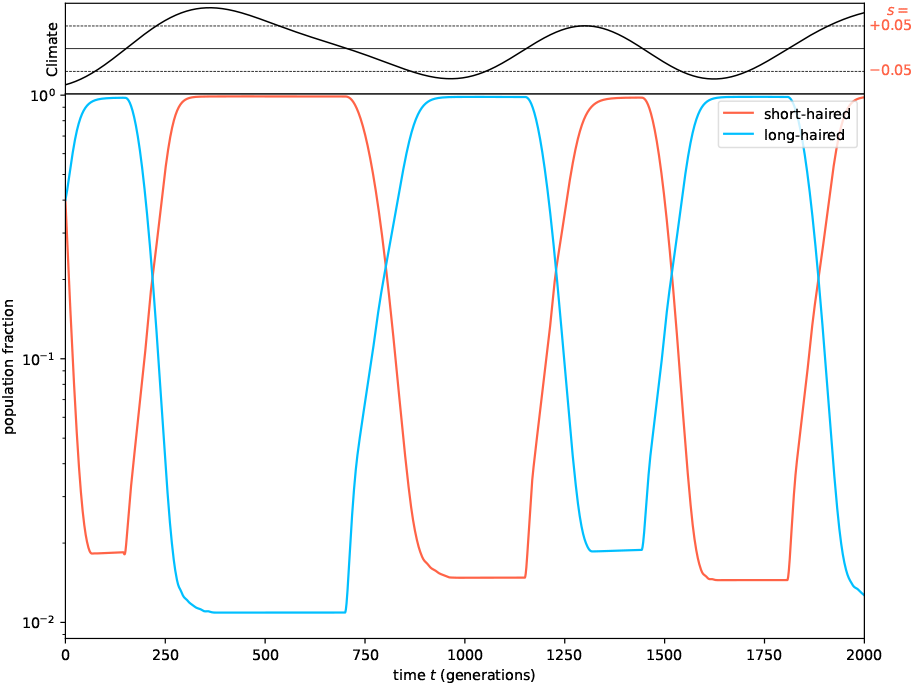
(Scenario 1.) Bet-hedging with slow environmental change. The favored phenotype increases to the environmental carrying capacity. The disfavored phenotype falls to a small equilibrium value, but never disappears, because it is the occasional offspring of the favored phenotype. When the environment reverses, the disfavored phenotype becomes favored.

Note the counter-intuitive result that, for small hedging *η*, the equilbrium is about 1*/s*_*∗*_ times larger than *η*, because the smaller disfavored population has (− *s*_*∗*_) losses proportional to its own size, but (*η*) gains proportional to the much larger favored population. The figure is a realization with *s*_*∗*_ = 0.05, *η* = 0.001, giving *f*_*u*_ ≈ 0.02, as seen. The fluctuations around this value are due to the stochastic nature of *G*(*t*).

We conclude that bet-hedging is quite a successful strategy, even with tiny amounts of diversity produced in each generation. The behavior remains qualitatively the same not only when *η* is small, but also in the opposite limit *η →* 0.5, with each generation an equal mixture of phenotypes. This might itself seem paradoxical, because, in equation (11), the loss of progeny to the favored phenotype far outweighs its selective advantage ∼ *s*_*∗*_. The resolution of the paradox is seen in the expression 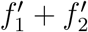, which is the net growth rate for the species (if not near carrying capacity). Equation (11) shows that, for this model, the *η*’s cancel exactly in this expression. In each generation the loss from one phenotype is exactly balanced by the gain from the other.

#### 3.2.2 Evolutionary Stability of Bet Hedging

Scenario 2, above, added the possibility that bet-hedging alleles might not be evolutionarily stable. The equations are,

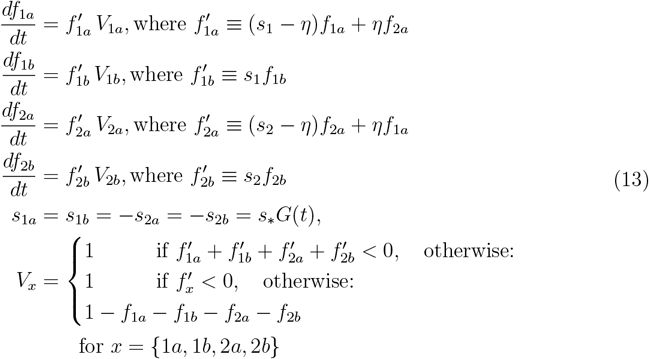

As compared to equation (11) (Scenario 1), equation(13) (Scenario 2) adds no new dimensionless parameters, so the two limits *T*_env_*/T*_fix_ ≶ 1 are generically again exhaustive (with checking on whether *η* does more than establishing the equilbrium of equation (12)). In the case *T*_env_*/T*_fix_ ≳ 1 (no figure shown), the wild type goes extinct just as in Scenario 0 (Figure 3), while the hedged variety has the same behavior as Scenario 1 (Figure 4).

The case of rapid environmental fluctuations, *T*_env_*/T*_fix_ ≲ 1, shown in Figure 5 is more interesting. We start the wild type with the population advantage. Both hedged and wild-type varieties are, by themselves, able to survive the rapid environment fluctuations. But, under adverse conditions the hedged variety loses less population, so it recovers more when the conditions change sign to favorable. The result is that it gradually, over many environment cycles, displaces the wild type—which goes extinct. Thus, in this model, bet hedging is evolutionarily stable against loss of the hedging phenotype.

**Figure 5:**
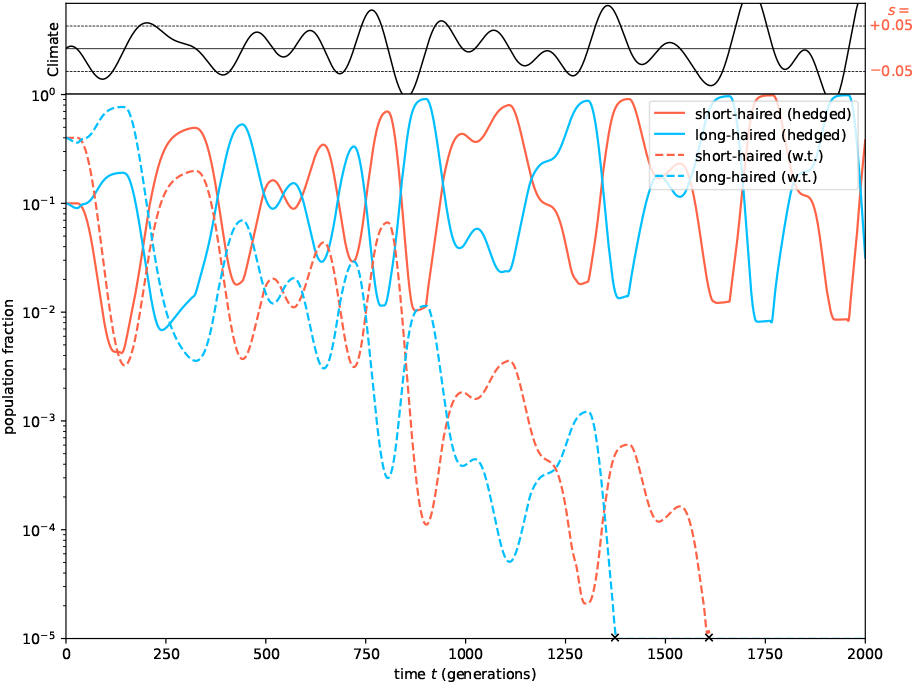
(Scenario 2.) Case of rapidly varying environment and with bet-hedging phenotypes (solid lines) competing against non-bet-hedging wild types (dotted lines). Although the wild phenotypes by themselves would survive indefinitely, they are slowly out-competed by the bet-hedging variety: In unfavorable times, the bet-hedged phenotypes suffer less loss, hence recover faster.

#### 3.2.2 Added Penalty for Hedging Trait

We saw in §3.2.1 that the introduction of a per-generation diversity parameter *η* produced no net loss of fitness for the species in the sense that the overall fitness 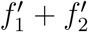 in equation (11) was unchanged. But what if the genomic pathway that creates the diversity itself imposes some additional loss of fitness *b >* 0? That is, the quantities 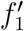 and 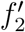 in equation (11) are replaced by

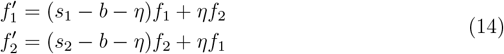

This in effect introduces a new dimensionless parameter, *b/s*_*∗*_, the burden *b* in relation to the intrinsic growth rate *s*_*∗*_. Obviously, the case *b/s*_*∗*_ ≳ 1 results in species extinction, because both long- and short-haired puffs will have contracting populations even in most favorable times. (We easily verify this by running the model, not here shown.)

Less obvious is whether a small but finite value *b/s*_*∗*_ ≪ 1 produces a secular population decline, versus the alternative of allowing complete recovery between environment cycles. This is addressed in Figure 6, which has *b/s*_*∗*_ = 0.2. The figure can be compared to Figure 4 (noting the change of scales). The salient differences are the “hooks” at the lowest populations of each cycle, e.g., at around *t* ≈ 600, 1250, 1800, and most notably 2600. The phenomenology is that the burden *b* drives a population loss of the unfavored species to below its diversity equilibrium, and the equilibrium is recovered only somewhat later, when the favored species fully recovers. But for *b/s*_*∗*_ ≪ 1, the equilibrium is indeed recovered.

**Figure 6:**
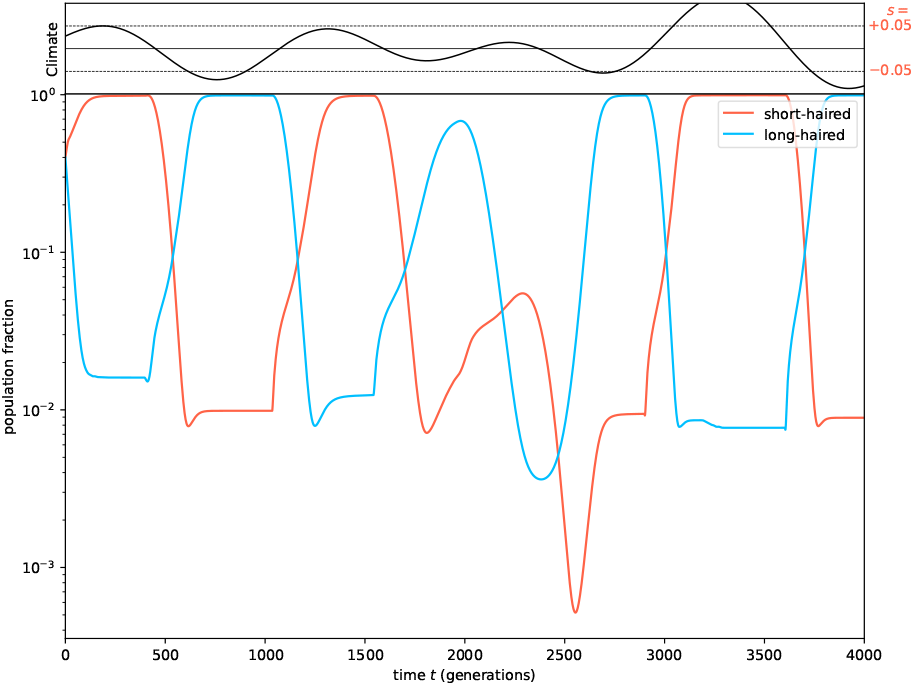
(Scenario 3.) Bet-hedging with additional penalty, case of slow environmental change. Here shown is the case where the penalty is not large compared to the intrinsic growth rate. The effect is to magnify initial phenotype declines (“hooks” in the figure). But there is time enough for recovery to the same equilibrium as in Figure 4.

As *b/s*_*∗*_ is increased towards ∼ 1, however the “hooks” rapidly deepen, reaching a values from which neither variety can recover. A realization of a borderline case with *b/s*_*∗*_ = 0.6 is shown in Figure 7. (This value of *b/s*_*∗*_ generally goes exinct in fewer cycles than this realization.)

**Figure 7:**
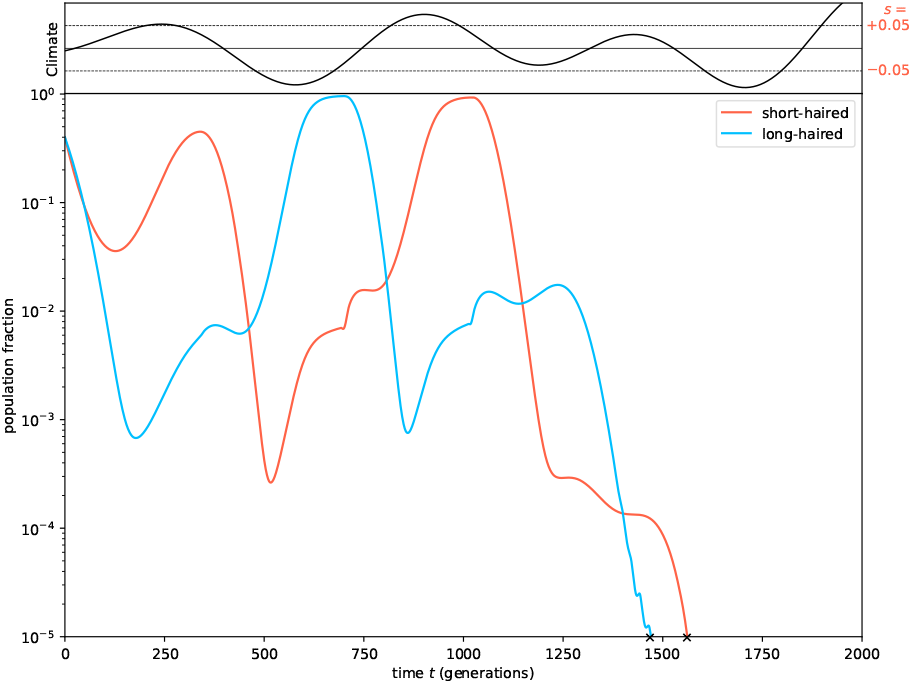
(Scenario 3.) Bet-hedging with additional penalty, case of slow environmental change. Here the penalty is large in comparison to the intrinsic growth rate. The disfavored phenotype does not fully recover after it becomes again favorable, leading to the eventual extinction of both phenotypes.

On the other hand, when *b/s*_*∗*_ ≪ 1 is truly satisfied, the realizations seen are practically indistinguishable from Figure 2 (when *T*_env_*/T*_fix_ ***≪*** 1) or Figure 4 (when *T*_env_*/T*_fix_ ≫ 1). That is, a small penalty *b* is tolerated with no long-term secular population decline. Nor, we find, does any tolerated level of burden *b* allow the wild-type (unhedged) population to invade.

### 3.3 Adaptive Mutation Models

We now put aside bet hedging as an adaptive strategy and turn to models in Scenario 4 (§2.2) “mutator strains”, that is, phenotypes with adaptively enhanced mutation rates from short-to long-haired, and vice versa. We assume Poisson random mutations occurring at a “effectual” rate (§2.2) of *µ* per individual per generation, implying a mean time between mutations of *N*_*∗*_*T*_mut_*/n*. (cf. equation7). These models evolve piecewise between mutation events by the equations (cf. equation 9),

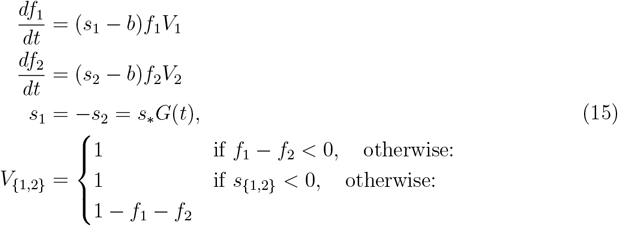

where now *b*, if nonzero, is an assumed “burden” due to deleterious mutations, namely, unfavorable mutations assumed to unavoidably accompany the useful ones. Mutation events are introduced as discontinuous step function increments of plus or minus a few times *s*_*∗*_. The exact small increment is irrelevant, since these mutations are already assumed effectual—a smaller value would simply move the events to slightly earlier. For simplicity we will assume equal rates of potentially rescuing mutations in the two directions, although this assumption could easily be relaxed.

There are now three dimensionless parameters: As before, *T*_env_*/T*_fix_ describes whether the environmental change is slow or fast compared with the fixation time of a favorable allele or extinction time of an unfavorable one. An additional new parameter is *T*_mut_*/T*_env_, which describes whether the environmental change is slow or fast compared with the time between favorable mutations. Finally the new parameter *b/s*_*∗*_ describes whether the burden *b* of other unfavorable mutations is small or large compared with the intrinsic growth rate *s*_*∗*_.

We first consider (§3.4) the survival prospects of a population with all mutator strains, and then (§3.5) look at whether such a population is susceptible to invasion by a strain that loses the enhanced mutability, and thus eliminates its burden *b*.

### 3.4 Survival of Mutator Strains

In equation (15), the effect of the burden parameter *b/s*_*∗*_ is easily disposed of. Simulations verify the obvious: For *b/s*_*∗*_ ≳ 1, the species goes rapidly extinct, because the effective intrinsic growth rate is negative most or all of the time. For *b/s*_*∗*_ ≲ 1, the effect of *b* is negligible. In the intermediate regime, *b* may tip the balance of the behavior due to the other parameters, but not by introducing any qualitatively new behaviors.

Among the remaining cases, consider first the case of slow environmental change, *T*_env_*/T*_fix_ ≳ 1, but different values of *T*_mut_*/T*_env_ (few to many mutations per environment time). Figure 8 shows an example with an insufficient mutation rate. While there are several mutations in each environmental epoch, these are (by definition) mutations to the disfavored phenotype (short- or long-haired), so are unable to fix in the population. The only exception is the mutation that occurs in the figure at *t* ≈ 700, which avoids extinction by occurring just when the environment is changing sign. That allows the short-haired phenotype to persist for one additional climate cycle. But that phenotype has no similar good luck in the timing of a mutation, so the species goes extinct.

**Figure 8:**
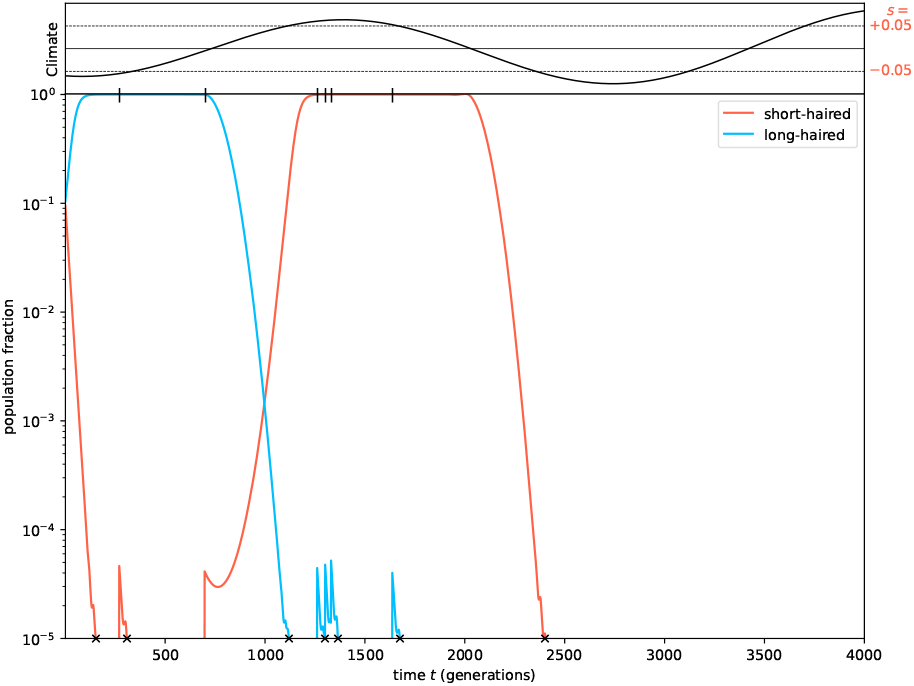
Mutator strain dynamics. Tick marks at the top of the lower plot, here and in subsequent figures, are where random mutations occur. Mutations seed the short- or long-haired phenotype that would in principle survive the next climate epoch, but if the mutations are too infrequent, each will die out before it becomes favored, as is the case in this realization.

Now compare Figure 9. Here the mutation rate has been increased to the point where there is (statistically, almost) always a mutation within what we here call the “fertile window”, defined as the time span *T*_*W*_ during which (i) the outgoing phenotype is still abundant enough to have reasonable probability of producing the mutation, and also (ii), the environmental sign change occurs before the thus mutated phenotype dies off.

**Figure 9:**
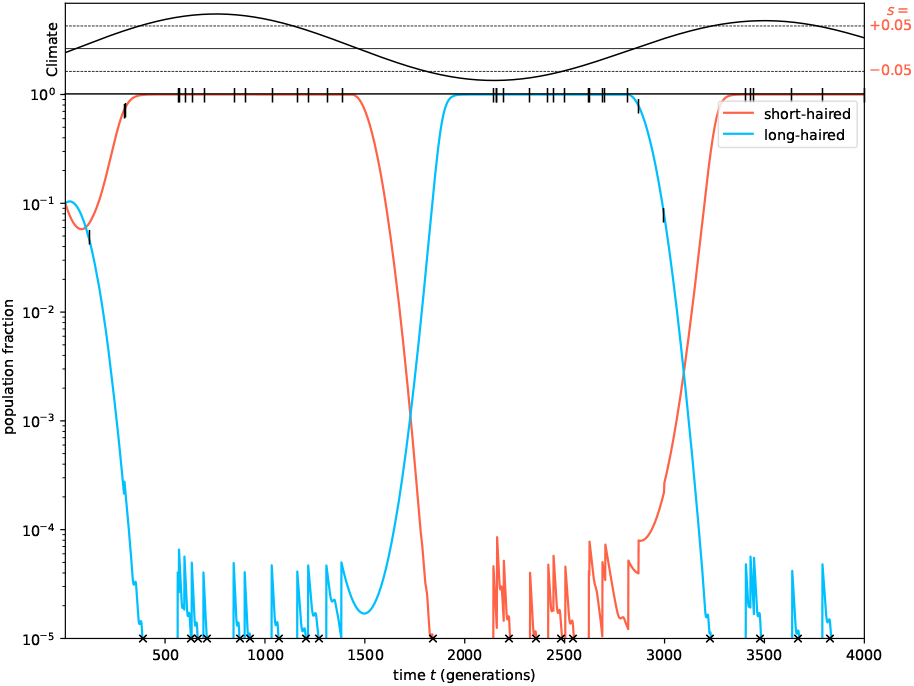
Like Figure 8, but now the mutation rate is large enough to seed the opposite phenotype in a “fertile window” close to a sign change in climate. See text for the calculation of that window size.

One might naively equate *T*_*W*_ ∼ *T*_fix_, on the grounds that *T*_fix_ is both the time during which the outgoing (now-unfavored) phenotype disappears and the now-favored phenotype grows to fixation. That argument, however, neglects the fact that *T*_fix_ is defined (equation 6) in terms of the mean or peak intrinsic growth rate *s*_*∗*_, while within*T*_*W*_ of a zero-crossing of *s*(*t*) the growth rate, positive or negative, is smaller by a typical factor *T*_*W*_ */T*_env_. (Here we assume that the environment changes continuously and not in discrete steps.) This slower rate implies a larger time window, so *W*_*T*_ obeys,

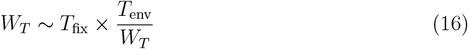

implying

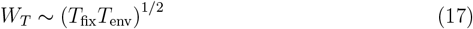

that is, the geometric mean of the fixation and environmental timescales, a perhaps unexpected result.

Regardless of mutation rate, both short- and long-haired phenotypes will survive in the regime of rapid environmental change, as was already seen in the baseline model of Figure 2, §3.1. That fact and, now, equation (17) allow a full characterization of the efficacy of a mutator strain in the parameter plane defined by *T*_env_*/T*_fix_ and *T*_mut_*/T*_fix_. This is shown in Figure 10. In the figure, it is important to note that the lines shown at definite values (e.g., 1) are in reality only typical within a factor *O*(1), and also that the lines are only fuzzy statistical boundaries: As a line is approached from one side, the behavior on the other side will start to appear in some statistical realizations.

**Figure 10:**
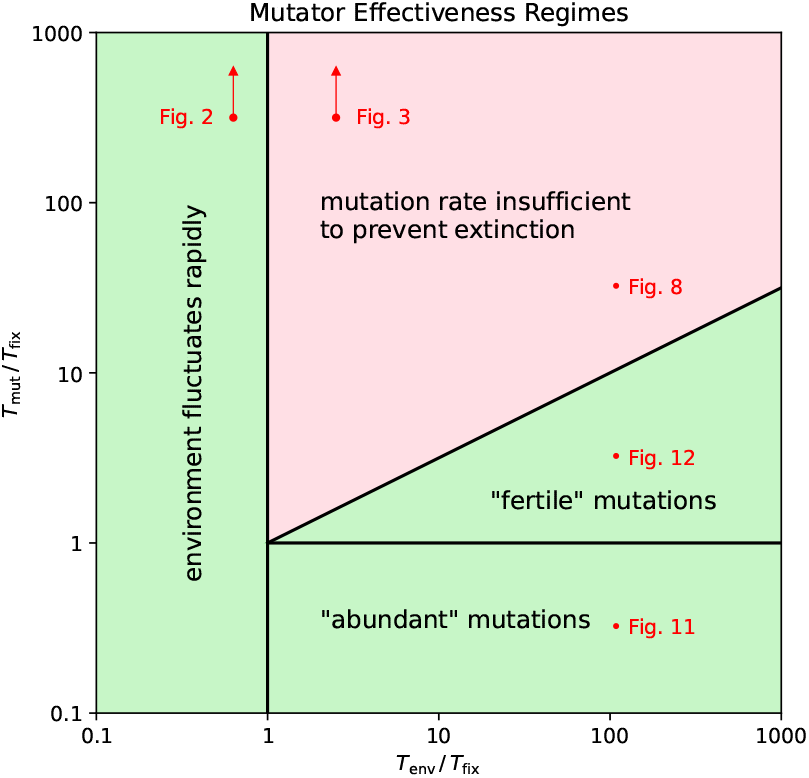
Regimes of mutator strain effectiveness as a function of the environmental timescale *T*_env_ and the mean wait time to the next mutation *T*_mut_, both normalized to the fixation timescale *T*_fix_. The species survives in the left-hand green rectangle because the environment changes rapidly enough to allow the survival of both phenotypes. It survives in the bottom green rectangle because favorable mutations occur more often then once per fixation (or extinction) time. In the green triangle, survival is due to a combination of just-frequent-enough mutations and just-slow-enough environmental change (see text for details.)

The non-intuitive feature of Figure 10 is the green triangle labeled “fertile mutations”. Here, potentially favorable mutations occur less often then one in *T*_fix_; but the environment is changing slowly enough to allow the broader fertile window, by equation (17).

Figures 2 and 3 in §3.1 can now be thought of as the limiting cases in Figure 10 of no mutations (i.e., *T*_mut_ → ∞). These are indicated in the figure. Figure 8, where the mutation rate is insufficient to prevent extinction, has its region in Figure 10 indicated. Similarly, Figures 11 and 12 complete the picture by showing a simulation in each of the remaining regions. (Figure 9, above, is approximately on the line between “abundant” and “fertile” mutations.)

**Figure 11:**
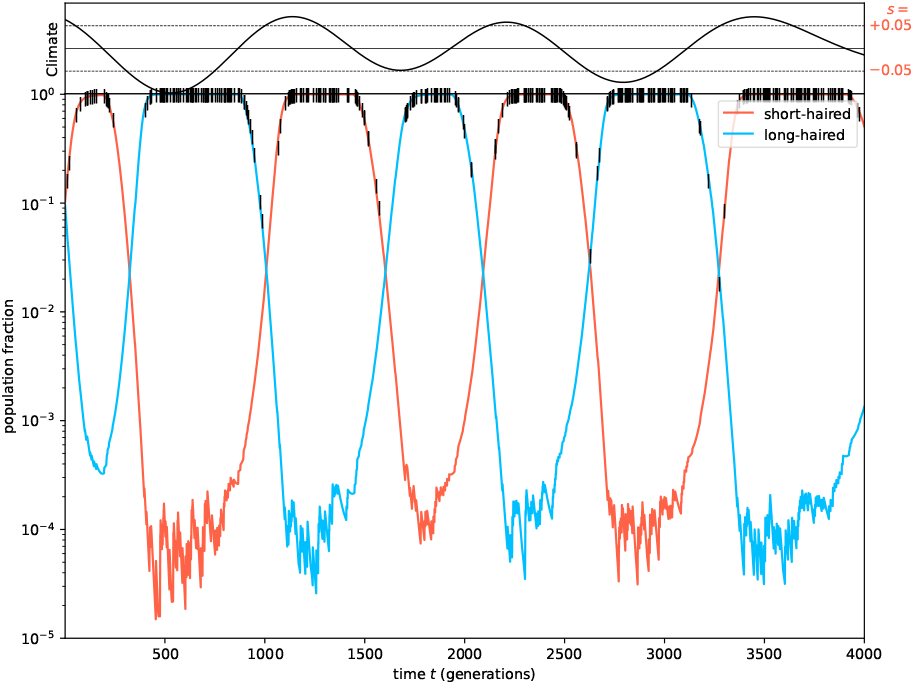
Typical simulation in the “abundant mutations” region of Figure 10. Several favorable mutations occur in every fixation time. The species survives regardless of the environmental change timescale.

**Figure 12:**
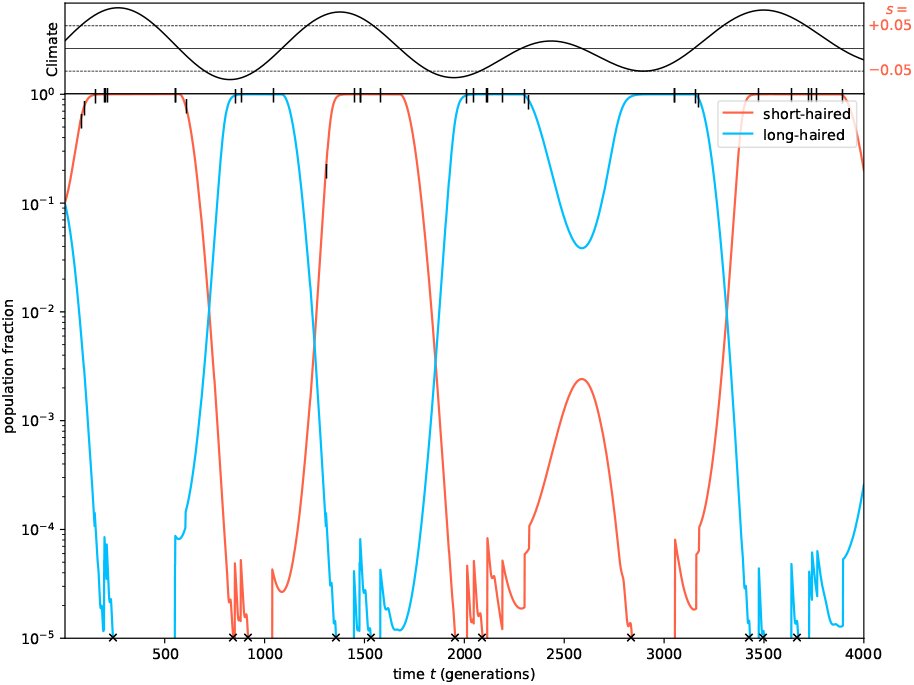
Typical simulation in the “fertile mutations” region of Figure 10. Fewer than one favorable mutation occurs per fixation time, but mutations are frequent enough to have several in a “fertile window” that depends on the relatively slow environmental change rate.

### 3.5 Non-Invasion of Mutators by Non-Mutators

In the model above, mutator phenotypes (both short- and long-haired) are penalized by a fraction *b/s*_*∗*_ of their intrinsic growth rate *s*_*∗*_. This suggests the possibility that a population of mutator phenotypes may be liable to invasion by a variant with loss of the enhanced mutation rate, shedding (we suppose) the penalty and reverting to wild type [23]. This invading population will by definition be unable to survive subsequent long-term environmental change.

The model equations (cf. equations 15 and 13) are piecewise

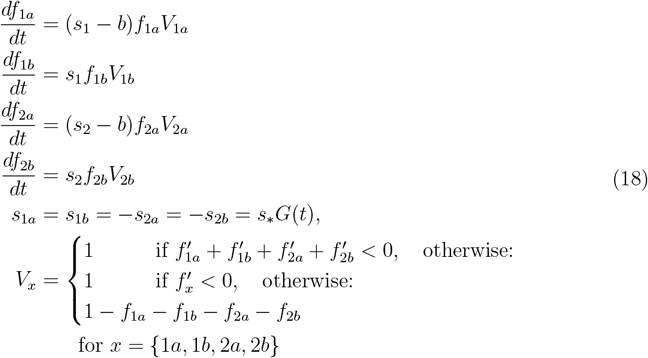

where subscript *a* denotes mutator strains, subscript *b* denotes wild types, and with discrete mutation events only between types 1*a* ↔ 2*a*.

A first analytic observation is that, no matter how long the environment is favorable to a variety (“1” or “2”), the wild type (“*b*”) can never completely replace the mutator (“*a*”). To understand this, consider simplified equations written as

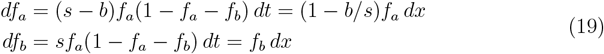

where

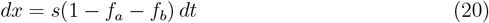

defines some (unknown but irrelevant) re-parameterization of time *t*. In terms of *x* and initial population fractions *f*_*a*0_ and *f*_*b*0_, the solutions are

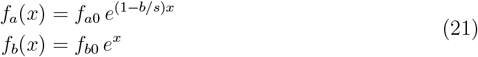

Since by equation (20) *dx →* 0 as *f*_*a*_ + *f*_*b*_ *→* 1, the asymptotic values *f*_*a∞*_ and *f*_*b∞*_ as *t* → ∞ are given by equation (21) with a value *x* = *x*_*∞*_ satisfying the transcendental equation

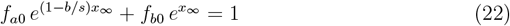

This equation is readily solved numerically for any values of *f*_*a*0_, *f*_*b*0_, and *b/s*. Then,

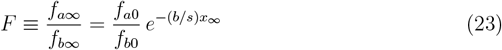

gives the final fraction of mutator strain relative to wild type, from which the final fraction of mutator strain relative to unity (the environmental carrying capacity) is readily calculated as *F/*(1 + *F*). Figure 13 shows this fraction for values *b/s* = 0.2 and 0.4, and for a range of starting fractions *f*_*a*0_ and *f*_*b*0_. The unshaded regions are where the invading wild type starts with a smaller prevalence than the mutator. One sees that over a wide range of initial conditions, the mutator equilibrates at *f*_*a*_ ≳ 0.1, which allows it to regain dominance on the next climate cycle.

**Figure 13:**
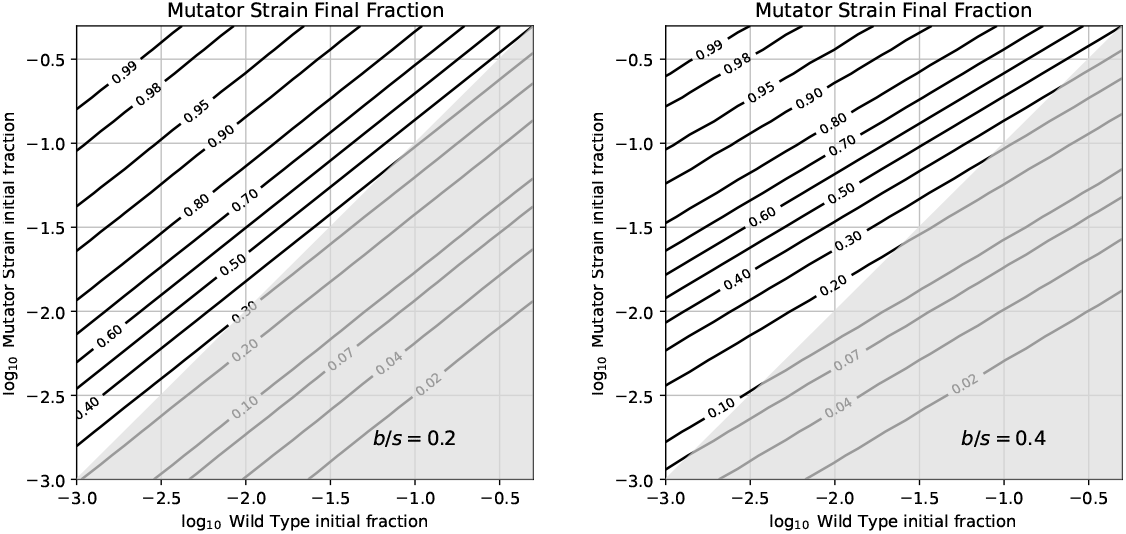
Despite a mutator strain’s assumed 20% (left) or 40% (right) penalty on intrinsic growth rate, it is able to resist invasion by a wild type of no penalty. The contour labels show the final equilibrium fraction of the mutator strain for a range of initial wild type fractions (abscissa) and mutator strain fractions (ordinate). A population initially dominated by mutators (upper left corner) remains so.

This behavior is shown in Figure 14, a numerical evolution of equation (18). Here a small initial seed population of long-haired wild type (dashed blue curve) succeeds in slightly dominating its mutator counterpart (solid blue curve) for one climate period, but goes extinct (at *t* ≈ 2300) after the climate changes sign. Meanwhile, the mutator has spawned frequent enough short-haired mutants (solid red curves) allowing the species to survive the sign change, as well as the one after that (at *t ≈* 3700).

**Figure 14:**
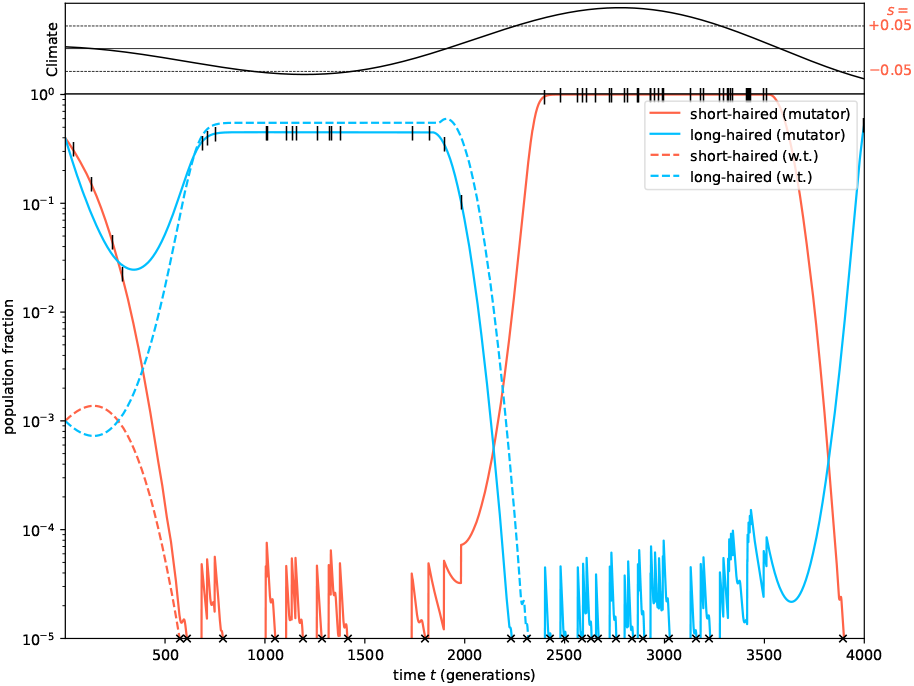
Attempted population invasion of a mutator population by a wild type with a larger intrinsic growth rate. The wild type (dashed curves) achieves partial dominance for one climate cycle, then goes extinct in the next. Meanwhile, the mutator (solid curves) comes to an equilibrium with a large enough population so as to spawn climate-resistant mutations that survive subsequent climate cycles.

## 4 Discussion

In the context of a unified set of population dynamical models and a stochastic but statistically well-defined model for long-term environment change (equations 2 and 3), we have simulated two possible genetic diversity mechanisms: bet-hedging, where a fraction of each generation’s offspring have characteristics adverse at present but useful in the future; and enhanced mutability, where mutations to that same (present-adverse, future-useful) phenotype are upregulated, but at the cost of other unfavorable mutations. While highly idealized (e.g., with only two phenotypes), the simulations allow outcomes to be identified and classified according to the values of a few non-dimensional parameters, namely the ratios of relevant timescales such as the fixation time, the environment-change timescale, and (for the mutator case) the average time to a relevant mutation.

Because evolution is blind, a population with one or the other mechanism for surviving climate variability might, in a favorable period, be out-competed by an invasion of the wild type. While short-term favorable, a successful such invasion would produce eventual species extinction when the climate changes; so we have tested, in the context of our models, whether such invasions succeed.

While many of the phenomena seen are unsurprising, a few deserve mention here:

1. Bet-hedging strategies can be successful even with very small diversity (fraction of offspring of the “wrong” phenotype per generation) *η*, in part because the equilibrium fraction of “wrong” phenotype is not *η* but a larger value ∼ *η/s*_*∗*_, where *s*_*∗*_ is the species intrinsic growth rate per generation (Figure 4 and equation 13).
2. A bet-hedging population is not successfully invaded by wild type. Either the two varieties co-exist at an equilibrium value until the next climate change, after which the wild type goes extinct; or else the bet-hedgers slowly out-compete the wild type over many climate cycles (Figure 2).
3. Bet hedging successfully tolerates small fractional penalties *b/s*_*∗*_ ≲ 1 to the species’ intrinsic growth rate *s*_*∗*_ without experiencing secular loss, but not large penalties *b/s*_*∗*_ ≳ 1.
4. Adaptive mutation, with a population of mutators, can succeed if the favorable mutation rate is as large in the population as a few per natural fixation time *T*_fix_, independent of the environmental change time *T*_env_.
5. Less obviously, it can also succeed with a smaller favorable mutation rate, but larger than a few per time (*T*_fix_*T*_env_)^1*/*2^ (equation 17 and Figure 10).
6. A mutator population, even one with a significant mutational burden *b/s*_*∗*_ is not readily invaded by a wild type without the burden, because the two reach a long-term equilibrium that allows the mutator to continue into the next climate cycle, when the wild type goes extinct (Figures 13 and 14).

We have not modeled how a population with the genetic diversity of our two phenotypes, the short- and long-haired puffs with either bet hedging or enhanced mutability, became established in the first place. As with any favorable trait, a chance occurrence is required. Coalescent theory [24] suggests that an effectuating mutation in a single individual, at just a right moment of environmental sign change, could have produced the common ancestor to all of the climate-change tolerant puffs that live on (fictitiously) today.

